# Vinegar and Its Active Component Acetic Acid Inhibit SARS-CoV-2 Infection *In Vitro* and *Ex Vivo*

**DOI:** 10.1101/2020.07.08.193193

**Authors:** Isabel Pagani, Silvia Ghezzi, Massimo Clementi, Guido Poli, Mario Bussi, Luca Pianta, Matteo Trimarchi, Elisa Vicenzi

**Author notes:** MT and EV share last authorship.

## Abstract

Effective and economical measures are needed to either prevent or inhibit the replication of SARS-CoV-2, the causative agent of COVID-19, in the upper respiratory tract. As fumigation of vinegar at low concentration (0.34%) ameliorated the symptoms of mild SARS-CoV-2 infection, we tested *in vitro* the potential antiviral activity of vinegar and of its active component, acetic acid. We here demonstrate that both vinegar and acetic acid indeed strongly inactivate SARS-CoV-2 infectivity in Vero cells. Furthermore, vinegar treatment caused a 90% inhibition of the infectious titer when directly applied to a nasopharyngeal swab transfer medium of a COVID-19 patient. These effects were potentiated if conduced at a temperature of 45 °C vs. 37 °C, a condition that is transiently generated in the upper respiratory tract during fumigation. Our findings are consistent and extend the results of studies performed in the early and mid-20^th^ century on the disinfectant capacity of organic acids and can provide an affordable home-made aid to prevent or contain SARS-CoV-2 infection of the upper respiratory tract.

## Introduction

At the end of 2019, a novel severe respiratory disease (coronavirus disease 2019, COVID-19) emerged in Wuhan, China and has since become pandemic in a few months, with *ca.* 11 million people infected worldwide as of today. COVID-19 is caused by a novel coronavirus called severe acute respiratory syndrome (SARS) CoV-2 to distinguish it from SARS-CoV that emerged in Guangdong province in China in 2003 and caused the severe clinical condition known as SARS. Like SARS-CoV, SARS-CoV-2 causes severe pneumonia that can lead to acute respiratory distress syndrome (ARDS) and death ^1^. However, unlike SARS-CoV, SARS-CoV-2 can causes on the one hand a multi-organ disease and hypercoagulation and, on the other hand, mild symptoms limited to the infection of the upper respiratory tract^2^. Indeed, high viral loads have been detected in the nasal swabs even in the presence of mild symptoms or in asymptomatic individuals ^3,4,5^.

Based on ancient medical tradition ^6^, in order to mitigate virus replication and its consequences in the upper respiratory tract mucosa, including acute anosmia (a frequent early symptom of SARS-CoV-2 infection ^7^), fumigation of vinegar diluted in boiling water at a concentration of 0.34% for 15 min has been adopted as an empirical practice in COVID-19 patients. Indeed, improvement of symptoms such as coughing and fever was observed in >90% of individuals ^8^.

In this regard, vinegar (derived from the biochemical processing of acetobacter species converting the ethanol of wine into acetic acid ^9^) has been used as a disinfectant for thousands of years and is commonly used to eliminate bacteria from fresh products ^10^ or as topical treatment for otitis externa ^11^. Acetic acid, the active component of vinegar, is also an effective disinfectant against mycobacterial infection ^12^.

Herein, we tested the potential antiviral activity of commercial vinegar and of its active component acetic acid against SARS-CoV-2 infection and replication *in vitro* and *ex vivo*.

## Materials and Methods

### SARS-CoV-2 plaque assay of *in vitro* and *ex vivo* virus replication

Vero cells were seeded at 2.5×10^5^ cell/well in 24-well plates in EMEM supplemented with 10% fetal calf serum (complete medium). Twenty-four h later, 50 plaque forming units (PFU) of a previously titrated SARS-CoV-2 isolate (GISAID accession ID: EPI_ISL_413489) were added to vinegar (from 0.28% to 0.008%) or acetic acid (from 0.1 to 0.5%) serially diluted (1:2) in PBS and incubated for 15 min at either 37 °C or 45 °C before addition to confluent Vero cells. Cell supernatants were discarded after 60 min and 1% methylcellulose (500 μl/well) dissolved in complete medium was added to each well. After 3 days, cells were fixed with formaldehyde/PBS solution (6%) and stained with crystal violet (1%; Sigma Chemical Corp.) in 70% methanol. Viral plaques were counted under a stereoscopic microscope (SMZ-1500, Nikon), as published ^13^.

A nasopharyngeal flocked swab (UTM® viral transport, COPAN Diagnostics Inc.) was obtained from a COVID-19 patient with severe symptoms. Fifty μl of the COPAN transport medium were diluted 1:2 with a vinegar solution at 0.28% either at 37°C or 45 °C for 15 min. The vinegar solution was then added to a monolayer of Vero cells that were seeded 24 h earlier at 2.5×10^5^ cell/well in 24-well plates in complete medium. As for in vitro infection, the supernatants were discarded after 60 min and 1% methylcellulose (500 μl/well) dissolved in complete medium was added to each well. After three days, plaques were stained and counted as described above.

All infection experiments were performed in a biosafety level-3 (BLS-3).

### Cell death detection assay

Ten μl samples of culture supernatant were transferred on a half black 96 well plate (Costar). To each well, 50 μl of the adenylate kinase detection reagent (ToxiLight^®^ BioAssay, Lonza) was added and the plate was incubated for 10 min at room temperature. Luminescence was measured in a Mithras LB940 Microplate Reader (Berthold Technologies). The results were expressed as Relative Light Unit (RLU).

### Statistical analysis

Prism GraphPad software v. 8.0 (www.graphpad.com) was used for all statistical analyses. Comparison among groups were performed using the one-way analysis of variance with the Bonferroni’s multiple comparison test.

## Results

### Vinegar and acetic acid inhibit SARS-CoV-2 replication in Vero cells

According to the concentration used for fumigation in SARS-CoV-2 infected patients ^8^, a concentration-response curve of viral inhibition by vinegar (diluted from 0.28% to 0.008% in PBS) was generated by a plaque assay previously optimized for Zika virus infection ^13^. Diluted vinegar effectively inhibited SARS-CoV-2 plaque formation by 80% at the top concentration of 0.28% with an inhibitory effective concentration 50 (IC_50_) of 0.08% (**Figure 1A**); of note is the fact that this vinegar dilution was not toxic to cells (**Figure 1B**) as revealed by the levels of adenylate kinase activity released in culture supernatants as an indicator of plasma membrane damage and cell necrosis ^13^. As the active component of vinegar is acetic acid, we next evaluated its anti-SARS-CoV-2 activity within the concentration range of vinegar. Indeed, incubation of SARS-CoV-2 with 0.5% acetic acid in PBS for 15 min at 37 °C reduced SARS-CoV-2 plaque formation by 89% with an IC_50_ of 0.14% (**Figure 1C**) without evidence of cytotoxicity (**Figure 1D**).

**Figure 1.**
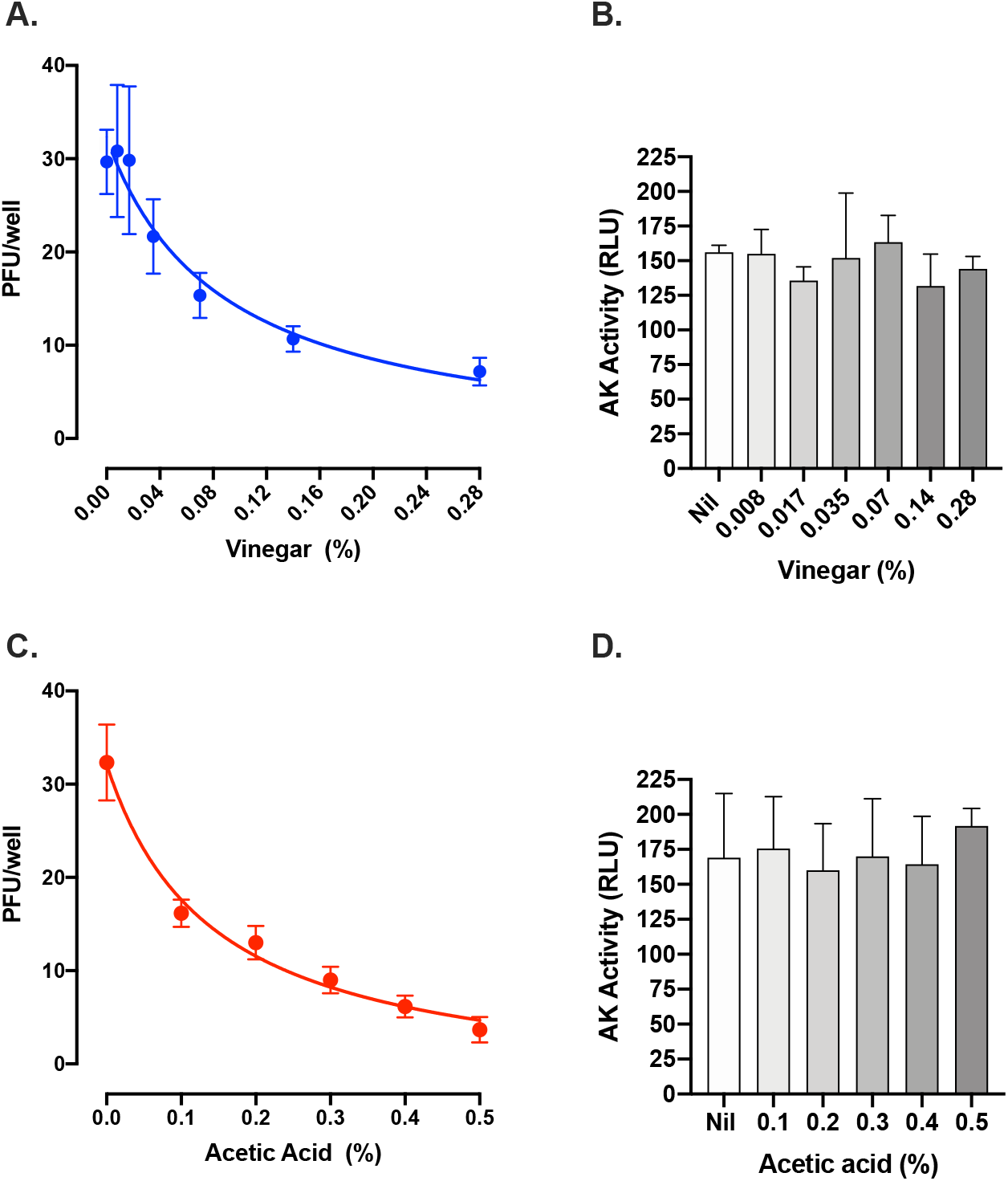
Vinegar and acetic acid, its active component, inactivate the infectivity of SARS-CoV-2. Inhibition of SARS-CoV-2 by commercial vinegar (**A**) and acetic acid (**C**) as measured by a plaque assay on Vero cells. **B.** Cell death after 48 h treatment with vinegar as measured by the activity of cell-associated adenylate kinase (AK) released in cell culture supernatants from Vero cells. Mean RLU (Relative Light Unit) ± SD of three independent experiments in duplicates are shown.

### Potentiation of vinegar and acetic acid antiviral effects by high temperature (t°)

As vinegar has been topically administered by fumigation in boiling water ^8^, an empirical determination of the t° was performed with a thermometer placed in the nostrils for 15 min during fumigation. The starting nose temperature was 54 °C and quickly decreased to 42 °C. Based on these empirical data, we tested the antiviral effect of vinegar diluted at 0.28% in PBS at 45 °C for 15 min. A significant decrease of plaque formation was observed by infecting Vero cells with virus treated at 45 °C as compared to untreated control at 37 °C (**Figure 2**). In addition, when the vinegar solution was heated at 45°C for 15 min, a significant additive reduction to 90% as compared to the virus inoculum treated at 37 °C was observed.

**Figure 2.**
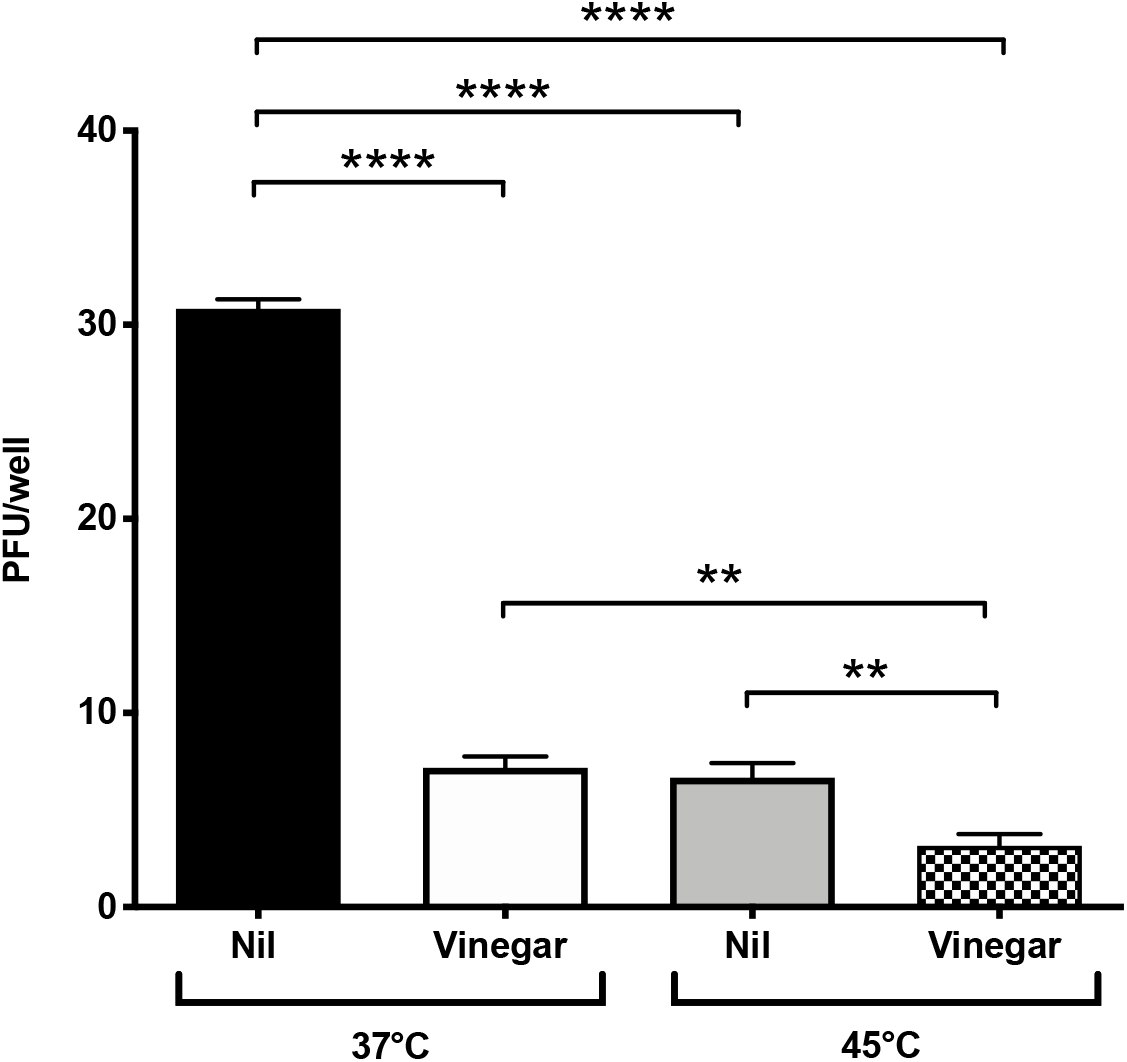
Temperature and vinegar inhibition activity on *in vitro* SARS-CoV-2 infectivity. Viral titers expressed as PFU/ml were obtained in a plaque assay of Vero cells infected with SARS-CoV-2 isolate that was treated with vinegar (0.28%) either at 37 °C or 45 °C, 15 min prior to addition to Vero cells. Nil means no treatment. Means ± SD of three independent experiments in duplicates are shown. ** indicate p<0.01, **** indicate p<0.0001 as calculated with one-way ANOVA with Bonferroni’s multiple comparisons test.

### Vinegar inhibits SARS-CoV-2 replication ex-vivo

We next evaluated whether vinegar could inactivate the infectivity of SARS-CoV-2 present in a nasopharyngeal swab transport medium. To this purpose, the flocked swab transport medium was directly tested in a plaque assay either in the presence of a vinegar solution at the concentration of 0.14%. When either 50 μl or 10 μl of transport medium were added to Vero cells, plaque formation proportional to the volume of transport medium was observed and vinegar significantly reduced the number of plaques **(Figure 3A)**. When tested at 45 °C, viral plaques generated by 10 μl of transport medium were reduced to almost zero (**Figure 3B**). As the plaque number were 5-fold less with 10 μl of transport medium than with 50 μl, the results were pooled as shown in **Figure 4**. The mean number of plaques in the control sample was 248±18 at 37 °C whereas at 45°C a significantly decreased (ca. 50%) number of plaques was observed. Vinegar reduced the plaque yield to *ca*. 50% at 37 °C and up to 90% at 45 °C.

**Figure 3.**
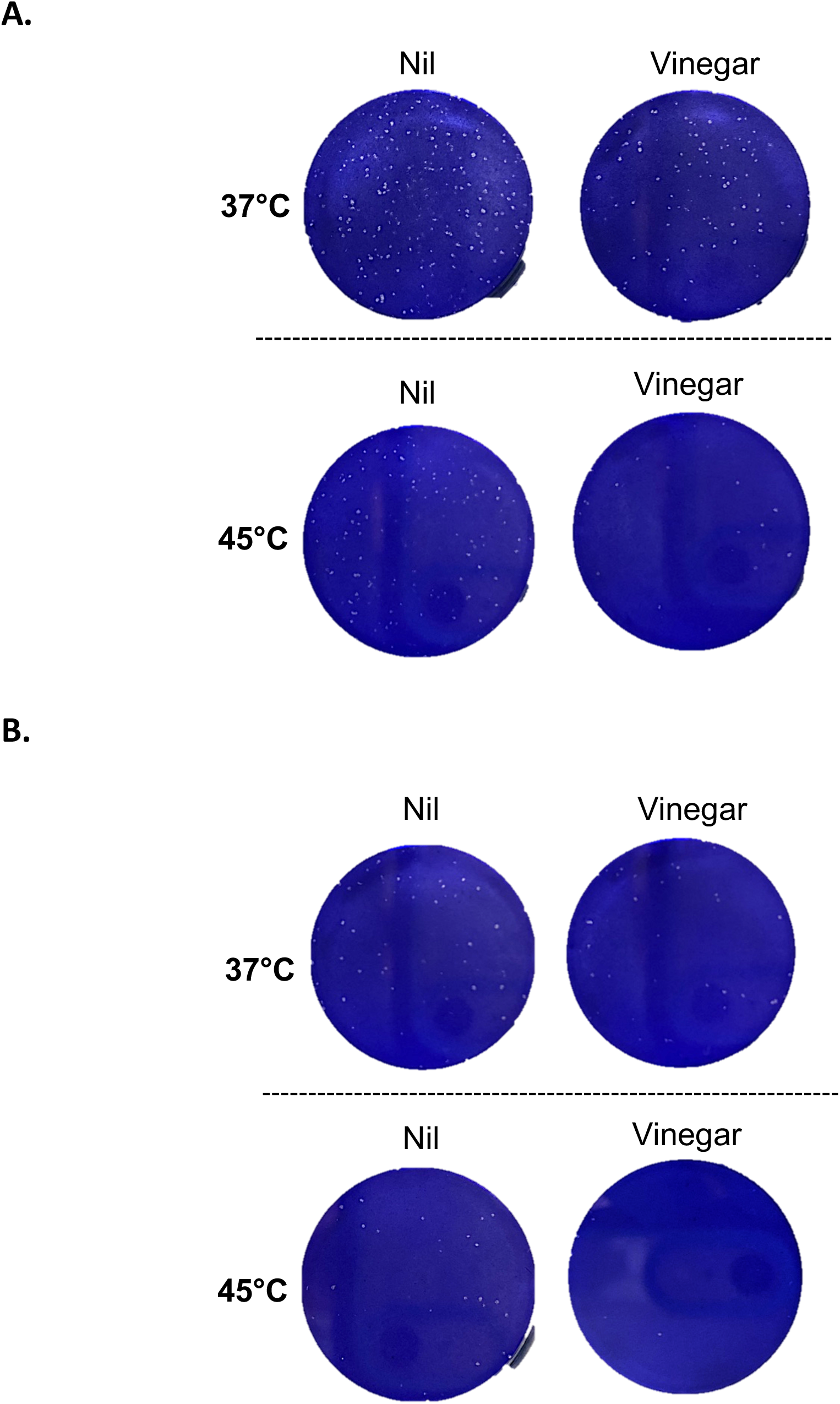
Temperature and vinegar inhibition activity on *ex vivo* SARS-CoV-2 infectivity. SARS-CoV-2 plaque formation was obtained by infecting Vero cells with 50 μl **(A)** and 10 μl **(B)** of a flocked swab transport medium from a COVID-19 patient. Inoculum was treated at 37 °C and 45 °C for 15 min in the presence or absence of vinegar (0.14%).

**Figure 4.**
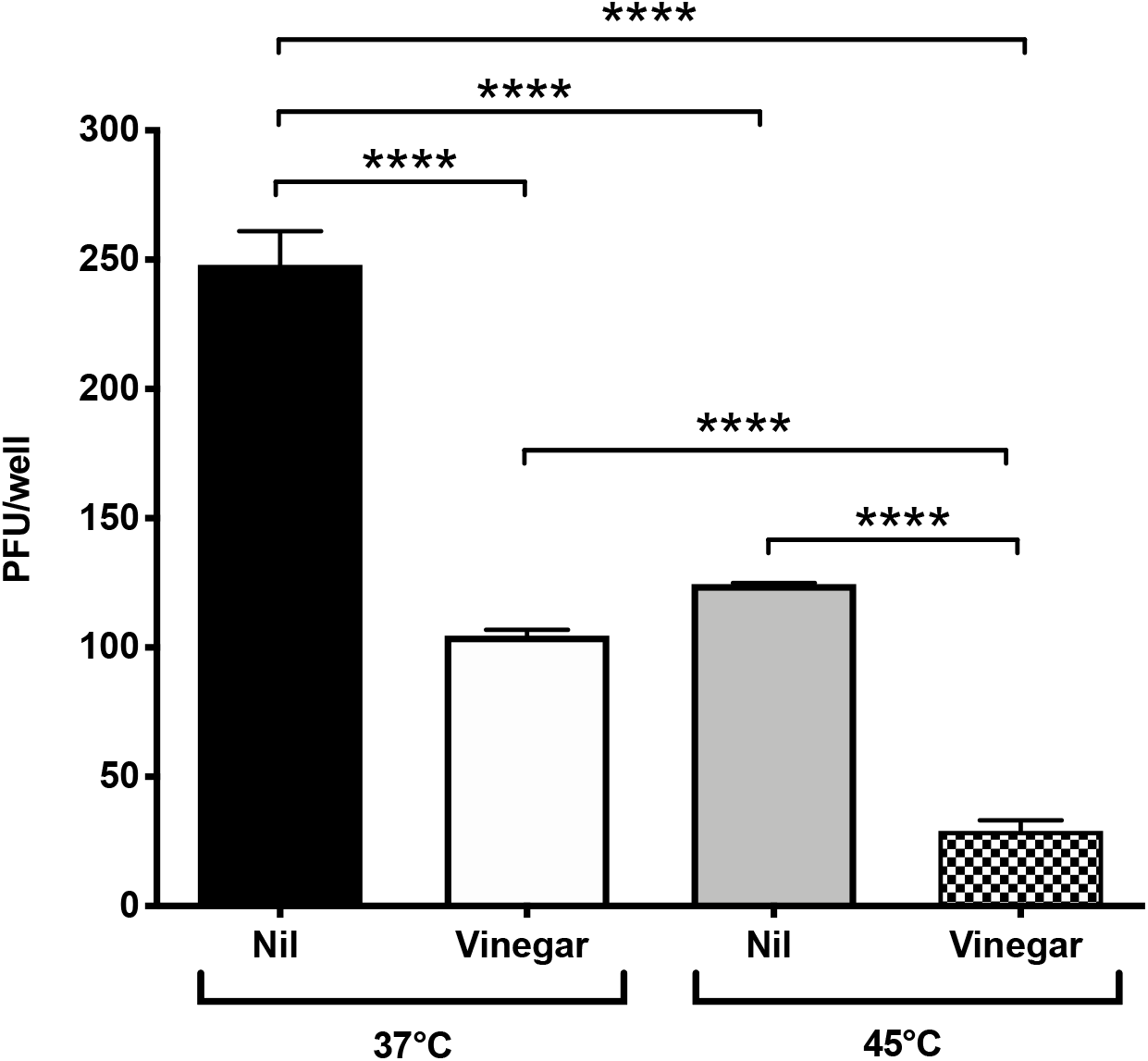
Temperature and vinegar additive effect on *ex vivo* SARS-CoV-2 inhibition. Viral titers expressed as PFU/ml were obtained in a plaque assay of Vero cells infected with the nasopharyngeal swab transport medium that was treated with vinegar (0.14%) either at 37 °C or 45 °C, 15 min prior to addition to Vero cells. Means ± SD of two independent replicates are shown. **** indicate p<0.0001 as calculated with one-way ANOVA with Bonferroni’s multiple comparisons test.

## Discussion

In the present study we demonstrate that both commercial vinegar and its active acetic acid component possess antiviral activity against SARS-CoV-2 in Vero cells infected with either a laboratory isolate or with the transport medium of a patient’s nasal swab in the absence of cell toxicity. Furthermore, when these experiments were conducted at 45 °C, an additive effect occurred in terms of antiviral activity in respect to what observed at 37 °C.

These experiments were conducted in Vero epithelial cells derived from an Africa Green monkey in 1957 and characterized by an absent production of interferon (IFN) in response to viral infection ^14,15^. For this reason, this cell line serves as preferential target of infection for several viruses, including influenza A virus ^16^, Zika virus ^13^, SARS-CoV ^17^ and SARS-CoV-2 ^18^. Thus, modulation of virus infection of Vero cells by any candidate antiviral agent cannot be attributed to indirect effect of an IFN response.

The concentration of acetic acid present in commercial vinegar varies from brand-to-brand and country to country. In Italy, it is typically found at 6-7% in vinegar derived from white wine, as used in this study, whereas in the United States it is usually 5%. Despite these differences, the IC_50_ as calculated from a non-linear fitting is relatively low (0.08%) and obtainable with vinegar sold at almost any percentage. As fumigation of vinegar has been used in mild COVID-19 patients in hot, boiling water, we here show that a temperature of 45°C had an additive effect in terms of inactivation of virus infectivity. Importantly, we here demonstrate that vinegar is effective in inhibiting the viral infectivity by lowering the viral titer present in a nasopharyngeal swab.

SARS-CoV-2 has been found to infect a wide range of cells of both the upper and lower respiratory tract expressing it entry receptor, i.e. angiotensin converting enzyme 2 (ACE2). Indeed, ACE2 expression analysis has shown that cells of the upper respiratory tract express higher levels of ACE2 as compared to those of the lower respiratory tract such as alveolar type II and type I pneumocytes ^19^. These differentially expression of ACE2 correlates with infectious titer of samples obtained from the upper respiratory tract than lower respiratory tract. Therefore, inhibition of viral invasion in the nasopharyngeal mucosa could prevent the invasion of the lower respiratory tract as previously observed in the case of antibiotic use during upper respiratory tract infection as a modality to prevent acute otitis media ^20^.

It has also been reported that, unlike SARS-CoV, the upper respiratory tract viral load can reach levels observed in the lower respiratory tract ^21^. Thus, inhibition of virus replication in the nasal mucosa by topic administration of diluted vinegar or acetic acid has the potential to prevent or curtail SARS-CoV-2 infection of the lower respiratory tract, thus reducing the occurrence of pneumonia and further clinical complications typically observed in COVID-19 patients ^1^.

As this simple treatment shows promising antiviral activity *in vitro* and amelioration of symptoms in mild SARS-CoV-2 infection, we suggest that this treatment has promising clinical use. While our preliminary studies warrant further testing in controlled clinical trials in patients with mild symptoms, in the absence of a fully effective antiviral therapy, this relatively inexpensive empirical treatment has the potential to mitigate the symptoms induced by SARS-CoV-2 infection of the upper respiratory tract by exerting local antiviral effects. These results also support the broader concept that compounds with anti-SARS-CoV-2 activity could be delivered as topical microbicides to inhibit of virus entry and replication in the upper respiratory tract. By reducing the viral load in the upper respiratory tract with a topical treatment, a significant impact in SARS-CoV-2 person-to-person transmission could also be achieved.

